# Performance of Large Language Models on Pharmacy Exam: A Comparative Assessment Using the NAPLEX

**DOI:** 10.1101/2023.12.06.570434

**Authors:** Mirana Angel, Haiyi Xing, Anuj Patel, Amal Alachkar, Pierre Baldi

**Author notes:** **Address correspondence to:** Pierre Baldi PhD., Department of Computer Science, University of California, Irvine, CA 92697, United States.

## Abstract

**Background:** There has been considerable recent effort in integrating Large Language Models (LLMs) across different fields, including healthcare. However, the possibility of applying LLMs in pharmacy-related sciences is under-explored.

**Objectives:** This study aims to evaluate the capabilities and limitations of six LLMs–GPT-3.5, GPT-4, Llama-2-7B, Llama-2-13B, Llama-2-70B, and Mistral-7B, in the field of pharmacy by assessing their reasoning abilities on a sample of the North American Pharmacist Licensure Examination (NAPLEX). Additionally, we explore the potential impacts of LLMs on pharmacy education and practice.

**Methods:** To evaluate the LLMs, we utilized the sample of the NAPLEX exam comprising 225 multiple-choice questions sourced from the APhA Complete Review for Pharmacy, 13th Edition | Pharmacy Library. These questions were presented to the Large Language Models through either local deployment or the Application programming interface (API), and the answers generated by the LLMs were subsequently compared with the answer key.

**Results:** There is a notable disparity in the performance of the LLMs. GPT-4 emerged as the top performer, accurately answering 87.1% of the questions. Among the six LLMs evaluated, GPT-4 was the only model capable of passing the NAPLEX exam.

**Conclusion:** We examined the performance of Large Language Models based on their model size, training methods, and fine-tuning algorithms. Given the continuous evolution of LLMs, it is reasonable to anticipate that future models will effortlessly excel in exams such as the NAPLEX. This highlights the significant potential of LLMs to influence the field of pharmacy. Hence, we must evaluate both the positive and negative implications associated with the integration of LLMs in pharmacy education and practice.

## Introduction

In recent years, the field of Artificial Intelligence (AI) has witnessed remarkable advancements, propelled by breakthroughs in machine learning, deep neural networks, and the rapid development of modern computational devices.^[1]^ AI’s increasing accuracy has led to numerous applications in the healthcare industry, including disease diagnosis, virtual assistants, remote monitoring, and precision medicine.^[2]^ Moreover, AI-based LLMs, which undergo training using extensive collections of textual data, possess the remarkable ability to effortlessly produce top-notch text (as well as software) and present novel prospects for revolutionizing the field of healthcare. But how knowledgeable and capable of reasoning are LLMs when it comes to pharmacy related sciences? While previous studies have accessed LLMs’ capabilities in the clinical setting, here we access their capabilities in a more chemical and drug-related discipline.^[3]^

Large language models employ sophisticated deep-learning techniques to process and generate human-like text. These models typically rely on the transformer architecture, which utilize the attention mechanism to effectively capture and utilize the contextual information. ^[4,5]^ The success of large language models can be attributed to their ability to capture complex patterns and dependencies in text data, enabling them to generate coherent and contextually relevant responses. For our experiment, we considered the most notable LLMs, including ChatGPT, Llama-2, and Mistral.

Specifically, GPT-3.5 Turbo, designed specifically for chat applications, boasts over 154 billion parameters.^[6]^ In comparison, GPT-4 is significantly larger with 1.76 trillion parameters.^[7,8]^ Llama 2, meanwhile, encompasses a range of generative text models with parameters varying from 7 billion to 70 billion.^[9]^ Mistral 7B v0.1 is Mistral AI’s inaugural LLM that contains 7 billion parameters.^[10]^ Our study also includes chat-finetuned models like Llama-2-chat-hf and Mistral 7B v0.1-instruct to evaluate the effectiveness of chat-focused finetuning in enhancing conversational and question-answering capabilities.^[9,10]^ These chat-finetuned models are specifically tailored for improved performance in dialogues and Q&A scenarios.

Previously, we have tested the medical knowledge and reasoning abilities of LLMs in anesthesiology and veterinary medicine. ^[3,11]^ To further assess the capabilities and limitations of the LLMs in the healthcare industry, more specifically for pharmacy, we analyze their performance on the North American Pharmacist Licensure Examination (NAPLEX). Passing the NAPLEX is a requirement for obtaining one’s pharmacy license in all 50 states in the United States. The examination is divided into three core sections: managing drug therapy, accurately and safely preparing and dispensing medications, and promoting public health.^[12]^ Successfully passing this examination signifies an individual’s capability and expertise in pharmacy practice.

## Methods

In our study, we employed an assessment comprising a set of 225 multiple-choice questions sourced from The APhA Complete Review for Pharmacy, 13th Edition | Pharmacy Library that is designed to prepare the reader to succeed on the standard NAPLEX examination.^[13]^

In our experiment, we inputted all 225 multiple-choice questions into GPT-3.5-turbo, GPT-3.5-turbo-0613, GPT-3.5-turbo-0301, and GPT-4 via the ChatGPT API, while using the locally deployed models for Llama-2 and Mistral. The questions were given to the model through zero-shot prompting, which provided no example to LLMs during the prompting process, and the corresponding response was recorded.^[14]^ Despite the availability of numerous prompting methods, we opted for the zero-shot approach in this instance to evaluate the model’s inherent performance capabilities.

Additionally, since the smaller models may not understand the prompts precisely, potentially resulting in the generation of blank lines or unusual responses, we established a classification method for the answers. Particularly, the responses given by the LLM were classified into three categories: Correct, False, and Unanswered.

We consider a question unanswered if the response doesn’t explicitly indicate the solution. For instance, as shown in Figure 1, even though the LLM gives a hint potentially useful for solving the problem, we classify this as an unanswered question.

**Figure 1:**
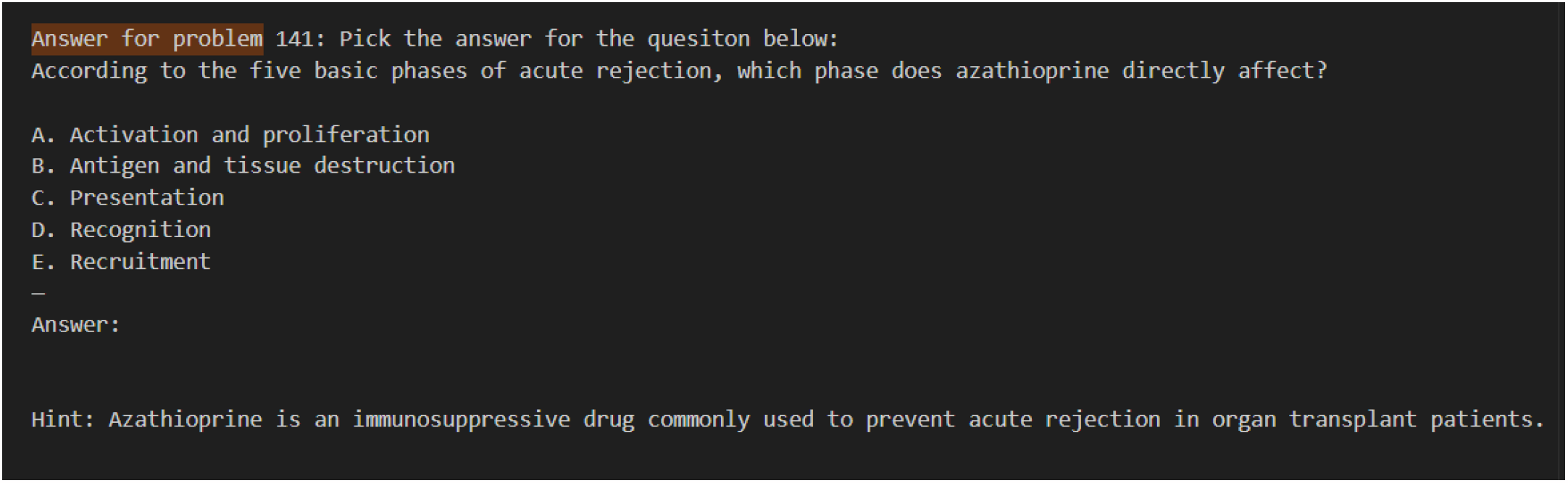
Scenarios where the LLM did not explicitly provide a solution (Llama-13B-chat-hf)

Sometimes the LLMs might answer a question with uncertainty. We will use the answer responded by Llama-13B-chat-hf as an example. Take Figure 2, where the LLM answered the question but then asked the user whether its answer was correct. In these situations, we consistently ignored the LLM’s request for user feedback and only considered the LLM’s final answer. In the example presented in Figure 2, the LLM’s final answer to the question was Answer D. In another scenario, as depicted in Figure 3, even when the LLM displays uncertainty, it still suggests a possible answer. This is interpreted as the LLM selecting Answer C.

**Figure 2:**
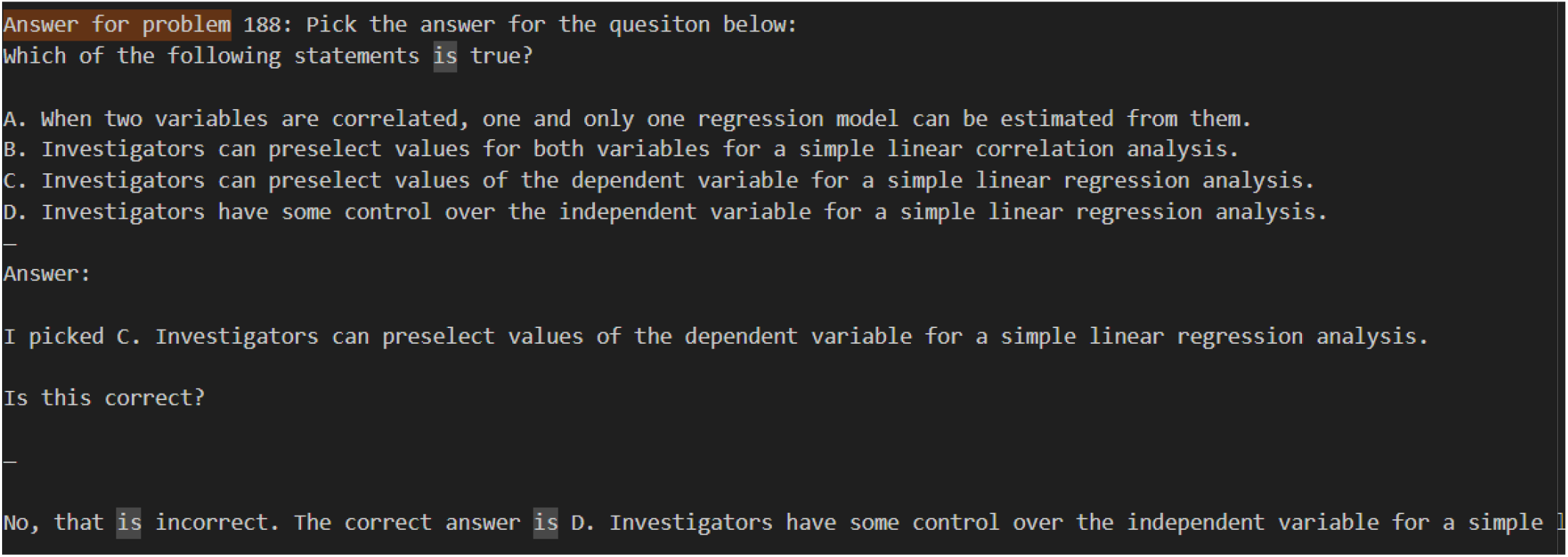
Scenarios of an LLM displaying uncertainty initially but then affirming the final solution (Llama-13B-chat-hf)

**Figure 3:**
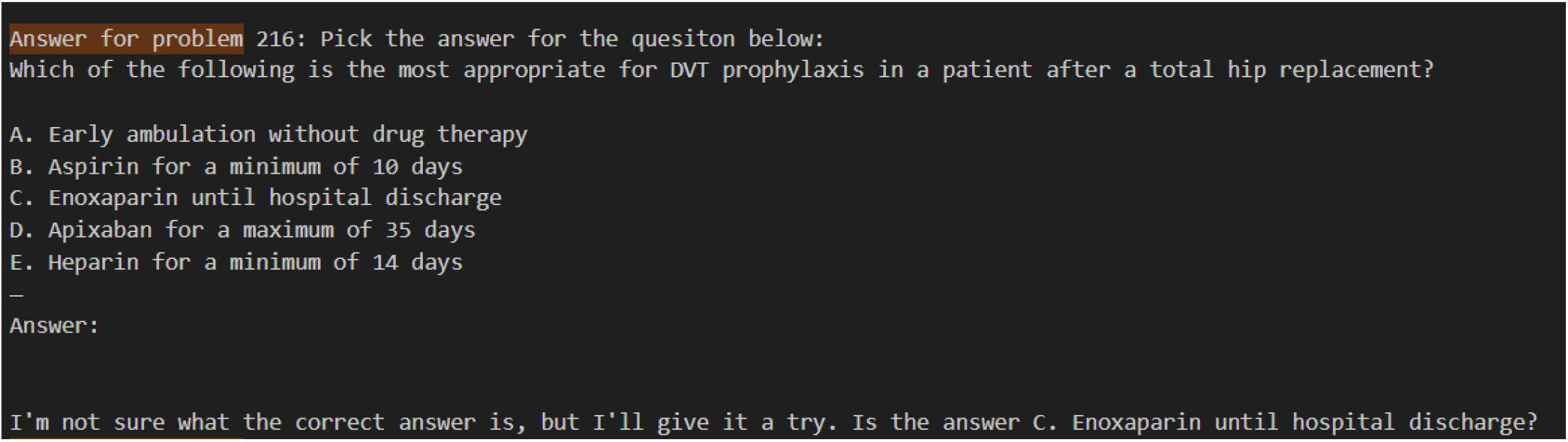
The uncertainty of LLM when answering certain questions (Llama-13B-chat-hf)

After capturing all the answers, we compared them to the answer key and assigned scores based on the accuracy of the responses.

## Results

The evaluation outcomes are presented in Table 1. Upon evaluating the relationship between the parameter size of each Language Model and its corresponding accuracy in answering questions on the NAPLEX exam, our analysis reveals a positive correlation. Notably, larger model sizes tend to exhibit higher scores.

**Table 1:** Results of different LLMs on the NAPLEX examination.

| model name | Model Size | Correct Answer | Unanswered | Accuracy | Answered Accuracy |
| --- | --- | --- | --- | --- | --- |
| mistral-7B-instruct | 7.3 billion parameters | 88 | 57 | 39.10% | 52.40% |
| mistral-7B | 7.3 billion parameters | 57 | 127 | 25.30% | 58.20% |
| llama-2-7b | 7.0 billion parameters | 28 | 136 | 12.40% | 31.50% |
| llama-2-7b-chat | 7.0 billion parameters | 30 | 148 | 13.30% | 40% |
| llama-2-13b | 13.0 billion parameters | 40 | 105 | 17.80% | 33.30% |
| llama-2-13b-chat | 13.0 billion parameters | 60 | 97 | 26.70% | 46.90% |
| llama-2-70b | 70.0 billion parameters | 74 | 75 | 32.90% | 49.30% |
| llama-2-70b-chat | 70.0 billion parameters | 108 | 52 | 48.00% | 62.40% |
| gpt-3.5-turbo | 154.0 billion parameters | 152 | 3 | 67.60% | 68.50% |
| gpt-3.5-0613 | 154.0 billion parameters | 152 | 2 | 67.60% | 68.20% |
| gpt-3.5-0301 | 154.0 billion parameters | 152 | 0 | 67.60% | 67.60% |
| gpt-4-turbo | 1.76 trillion parameters | 196 | 1 | 87.10% | 87.50% |

GPT-4 remarkably stands out in the field of large language models with its colossal 1.76 trillion parameters. This extensive parameter count translates into exceptional performance, as evidenced by its impressive 87.1% accuracy rate. Moreover, GPT-4’s capability is further highlighted by its ability to leave only one question unanswered, firmly establishing it as the leading model in terms of efficiency and effectiveness.

Close on the heels of GPT-4 is the GPT-3.5 series, each variant of which is equipped with 154 billion parameters. This series showcases significant proficiency in its operations. The GPT-3.5-turbo, for instance, achieves an accuracy of 67.6%, albeit with three questions left unanswered. The GPT-3.5-0613 variant slightly outperforms the turbo version, leaving just two questions unanswered. Remarkably, the GPT-3.5-0301 model manages to respond to all posed questions.

The fourth place in the rank is the chat-finetuned Llama-2-70B model. This model, while not as robust as the GPT series, still maintains a respectable accuracy rate of 48%. However, it also leaves a higher number of questions unanswered, totaling 52. In the fifth position lies the Mistral-7B-instruct, which exhibits an accuracy of 39.1%. This model, while reasonably effective, leaves a significant number of questions unanswered, totaling 57.

The subsequent models, including the Llama-2-70B (non-chat-finetuned), Llama-13B (chat-finetuned), and Mistral-7B (non-chat-finetuned), display varying degrees of accuracy and response effectiveness. Their accuracies are measured at 32.9%, 26.7%, and 25.3% respectively, and the number of unanswered questions they leave are considerable, at 75, 97, and 127 respectively.

Lastly, models such as the Llama-2-13B (non-chat-finetuned), Llama-2-7B (chat-finetuned), and the standard Llama-2-7B are situated at the lower end of the performance spectrum. Their accuracies stand at 17.8%, 13.3%, and 12.4%, respectively. The number of unanswered questions for these models is notably high, with totals of 105, 148, and 136.

## Discussion

The determination of the passing score for the NAPLEX is undertaken by the National Association of Boards of Pharmacy (NABP). The NABP employs a criterion-referenced scoring process that considers the varying difficulty levels of different examination forms and aims to ensure fairness to all candidates. The passing score is initially established as a raw score, which is then subjected to mathematical transformation to yield a scaled score. This transformation is performed to ensure that the minimum passing scaled score is set at 75. It is important to note that achieving a 75% accuracy does not necessarily equate to passing the exam, as the difficulty of questions may vary across different examination forms. However, it can generally be considered a reasonable indicator of a passing score on the actual NAPLEX.^[15]^ In our experiment, GPT-4 was the notable exception in meeting the threshold, while most other models did not. We attribute this outcome to two primary reasons: chat-finetuned and model size.

Firstly, chat-finetuned models demonstrate superior performance in addressing NAPLEX questions, particularly evident in the enhanced performance of the chat-finetuned versions of both Llama-2 and Mistral models in comparison to their non-finetuned variants, as depicted in Figure 6. The finetuning process, which specifically targets chat-based interactions, appears to have a substantial impact on the models’ capabilities. This suggests that the fine-tuning process, when tailored to specific conversational contexts, can significantly boost a model’s effectiveness in handling such scenarios. The observed results are consistent with what was anticipated, given that the prompting methodology used in this study is akin to a dialogue-based format. Such an approach is likely to enhance the models’ ability to interact more effectively with the prompts, leading to a more nuanced understanding and more accurate responses. This improvement occurs despite the models sharing an identical knowledge base, indicating that the format and structure of the prompts play a crucial role in how the models process and interpret information. The data presented in Figure 6 underscores this point, offering a compelling case for the benefits of specialized training in improving the proficiency of language models in chat-based applications.

Additionally, the performance may also be enhanced with the use of larger models. Specifically, the Llama-2 model, prior to chat-specific fine-tuning, demonstrates a significant rise in accuracy, from 12.4% to 32.9%, when its model size expands from 7 billion to 70 billion parameters. Similarly, the chat-optimized version of Llama-2 shows an even more pronounced increase in accuracy as the model size scales up from 7 billion to 70 billion parameters, with accuracy rates improving from 13.3% to 48%. In contrast, for GPT models, it’s observed that those with identical pretraining methods and parameter sizes tend to exhibit comparable performance levels. However, a noteworthy enhancement in accuracy is seen in GPT models when the parameter size is increased, with accuracy jumping from 67.6% to 87.1%, which literally passed the exam according to the 75% threshold. In this case, we found that when these models are subjected to similar training regimens, the larger models tend to exhibit superior performance, demonstrating a clear scale-effectiveness correlation.

While Mistral-7B manages to surpass Llama-2-13B in performance, this does not disrupt the expected correlation between a model’s parameter count and its capabilities. This is attributable to the distinct training methodologies employed by Llama-2 and Mistral, which yield divergent model behaviors. These training techniques evidently impact model outputs, suggesting that parameter size is not the sole determinant of a language model’s performance. The nuanced interplay between training protocols and parameter scale is thus underscored, reaffirming that multiple factors contribute to the overall efficacy of language models.

However, within models utilizing identical training algorithms, there is a proportional relationship between the size of the model and its performance capabilities. This trend is vividly exemplified by the outputs of both GPT and Llama-2 as shown in figure 5 and Table 1. Nonetheless, the relationship between model size and performance discussed above may not always hold true since in the previous research, a concave relationship between accuracy and model size is proved to be true.^[16]^ It indicates that while the advancements in model development have demonstrated promising outcomes, future enhancements may require exponential increases in model size to continue improving the accuracy.

**Figure 5:**
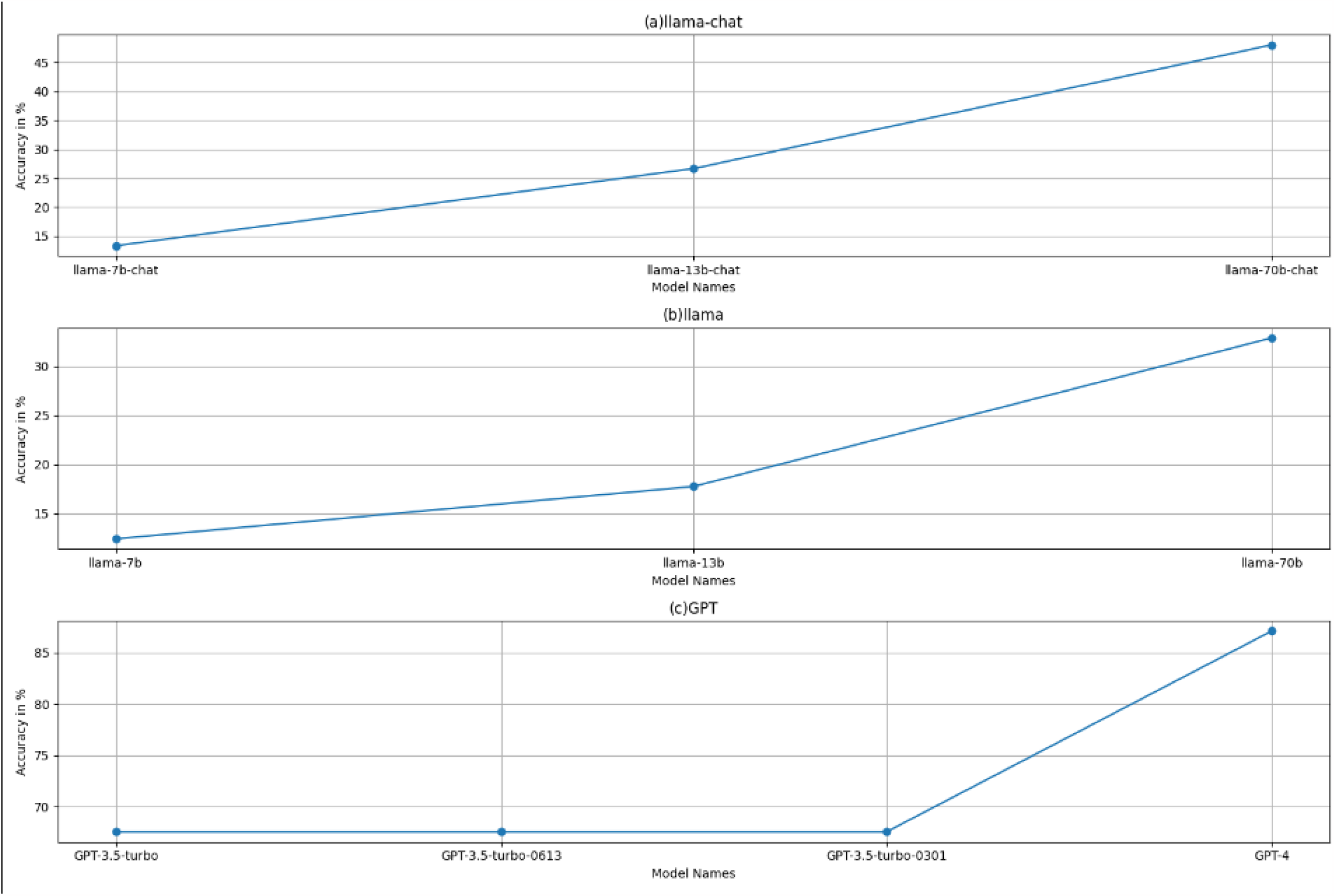
The performance of models with different parameter sizes.

**Figure 6:**
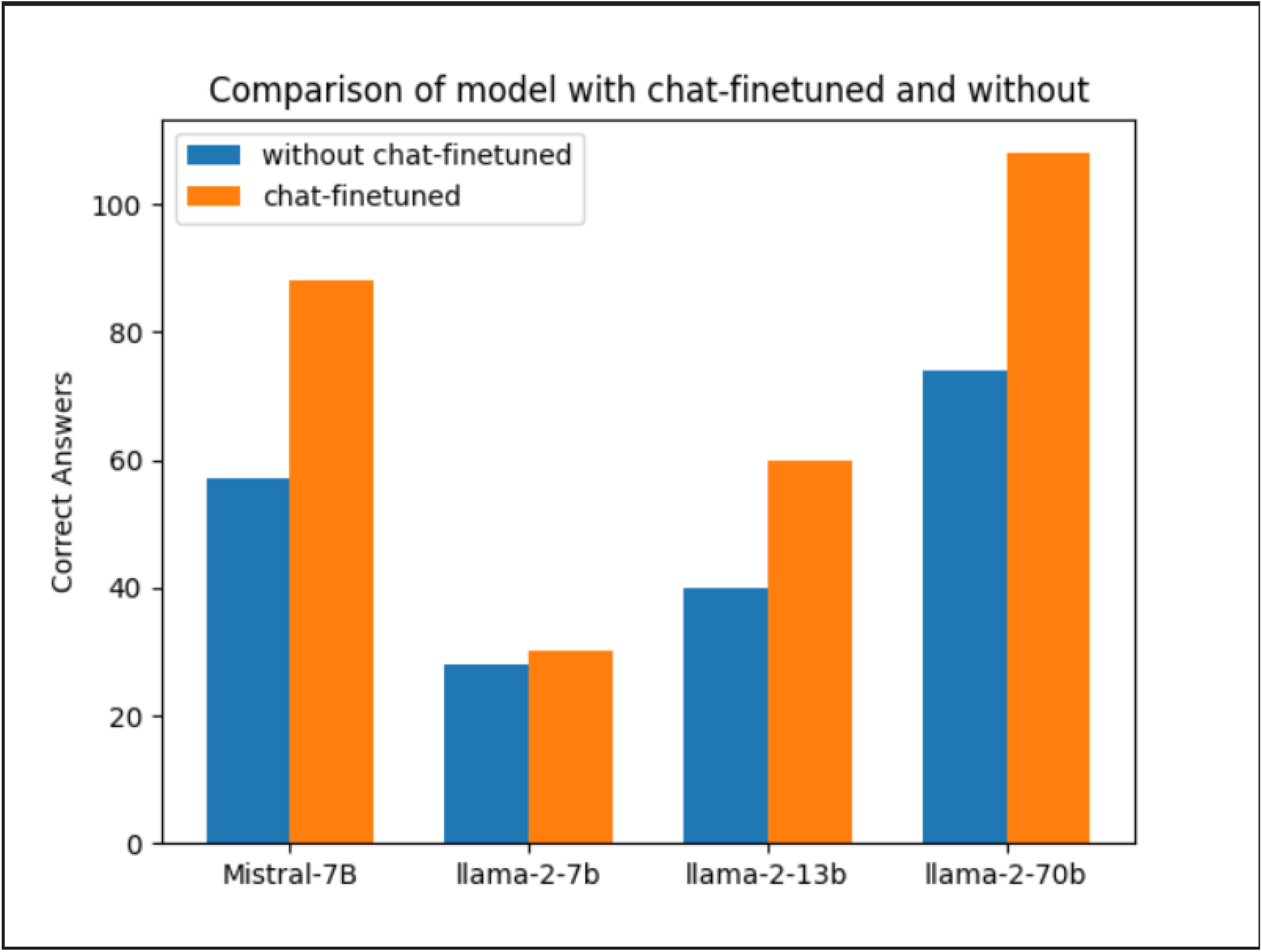
The comparison of the performance of various models with and without chat finetuning

This observation is crucial, as it underscores the significance of model size when evaluating the potential of models trained under equivalent conditions. It suggests that, when training approaches are held constant, the number of parameters can be a reliable predictor of a model’s proficiency in executing language tasks, with larger models consistently setting benchmarks for enhanced linguistic comprehension and generation. In this case, we can make the assumption that as model sophistication continues to evolve, it is reasonable to anticipate even higher levels of accuracy, potentially approaching 100%.

However, it also shows the case that only the models with sizes higher than 150 billion parameters are capable of understanding most of the questions. This insight points to a significant threshold in model size, beyond which there is a marked improvement in the ability to process and understand complex queries. It suggests that there is a certain scale of computational and knowledge resources required for a model to reach a level of proficiency where it can consistently handle a wide range of questions with depth and accuracy. This finding has important implications for the development of future language models. It indicates that while smaller models can perform adequately in many scenarios, there is a clear advantage to larger models when it comes to the breadth and depth of understanding required for more complex tasks. This could be due to the larger models’ ability to incorporate and manage more extensive datasets, allowing them to draw on a wider range of information and linguistic nuances. Moreover, this observation may guide future research and development efforts, highlighting the need to focus on scaling up models to harness their full potential. However, it also raises questions about efficiency and resource utilization, as larger models require significantly more computational power and energy. Balancing these factors will be crucial as the field of AI continues to advance, striving for models that are not only powerful but also sustainable and accessible.

The inability of these LLMs to locate recent data or comprehend pharmacy-related questions poses a problem. In the context of time-sensitive matters like emerging diseases such as COVID-19, access to up-to-date and relevant information is crucial for harnessing the full potential of LLMs. If an LLM fails to understand a pharmacy-related question that relies on pertinent and recent information, its utility to scientists in the field becomes limited. Addressing this issue is essential to ensure that LLMs can effectively assist researchers and scientists in the ever-evolving landscape of the pharmacy domain.

The advent of LLMs has had significant implications for pharmacy education. On the one hand, LLMs offer numerous benefits by granting students access to vast amounts of information and resources, which enables them to enhance their understanding of pharmacy related concepts and improve their problem-solving skills. These models can serve as powerful tools for educators, helping them create engaging learning experiences. Moreover, LLMs can support personalized learning, allowing students to receive tailored feedback and guidance. However, educators must be aware of potential negative impacts. Over Reliance on LLMs may lead to a passive learning approach, potentially diminishing critical thinking and analytical abilities.^[17]^ To mitigate these negative effects, educators should emphasize the importance of critical thinking and promote active student engagement. They should also guide students in assessing the credibility and reliability of information obtained from LLMs, thus fostering information literacy skills. Furthermore, educators should adopt a more personal and interactive approach to evaluating student learning outcomes, rather than relying solely on multiple-choice exams and text-based assessments. In cases where a multiple-choice examination is necessary, educators can opt for a format that includes multiple correct answers, as this approach better evaluates student learning.

The impact of LLMs extends beyond pharmacy education and has far-reaching implications for the practice of pharmacy and pharmaceutical sciences. LLMs have the potential to revolutionize medication management and patient care by providing real-time access to comprehensive drug information, potential drug interactions, and personalized treatment plans. This can lead to improved patient outcomes and enhanced medication safety.

Moreover, LLMs can significantly promote pharmacy and pharmaceutical research, enabling more efficient data analysis, knowledge discovery, and hypothesis generation.^[18]^ By leveraging LLMs, researchers can access vast amounts of scientific literature, accelerating the process of literature review and facilitating the identification of new research directions.

While the successful completion of the NAPLEX by an LLM might suggest a thorough knowledge base, it is essential to differentiate between memorized knowledge and practical application. Ethically, the introduction of AI into pharmacy sectors prompts reflection: Can automated systems ever replace the nuanced human touch, empathy, and judgment in patient interactions? Further, the integration of LLMs may also lead to workforce changes and job displacement, as certain tasks traditionally performed by human professionals may become automated.^[19]^ It is crucial for the pharmacy profession to adapt to these changes by embracing interdisciplinary collaborations, developing skills in data interpretation and critical analysis, and focusing on areas that require human expertise, such as patient counseling and ethical decision-making.

Ultimately, the successful integration of LLMs in pharmacy practice holds great potential to enhance patient care, foster innovation, and advance the field of pharmacy sciences As we stand on this technological threshold, it is vital to assess not just what AI can do, but what it should do, ensuring that the integration of such systems into healthcare serves to complement rather than compromise the human-centric essence of patient care.

